# Analysis of differential expression of matrix metalloproteins and defensins in the nasopharyngeal milieu of mild and severe COVID-19 cases

**DOI:** 10.1101/2024.05.12.593784

**Authors:** Khekashan Imtiaz, Nida Farooqui, Khalid Ahmed, Alnara Zhamalbekova, Muhammad Faraz Anwar, Khitab Gul, Azhar Hussain, Antonio Sarría-Santamera, Syed Hani Abidi

**Affiliations:** Department of Pathology and Laboratory Medicine, Aga Khan University, Karachi, Pakistan; Department of Biological and Biomedical Sciences, Aga Khan University, Karachi, Pakistan; Department of Biochemistry, Bahria University Medical and Dental College, Karachi, Pakistan; Department of Biosciences, Muhammad Ali Jinnah University, Karachi, Pakistan; Department of Biomedical Sciences, Nazarbayev University School of Medicine, Astana, Kazakhstan

**Keywords:** COVID-19, SARS-CoV-2, matrix metalloproteinases, defensins, COVID-19 mild and severity disease, biomarkers

## Abstract

**Introduction:** A subset of COVID-19 disease patients suffers a severe form of the illness, however, underlying early pathophysiological mechanisms associated with the severe form of COVID-19 disease remain to be fully understood. Several studies showed the association of COVID-19 disease severity with the changes in the expression profile of various matrix metalloproteinases (MMPs) and defensins. However, the link between the changes in the expression of matrix metalloproteinase (MMPs) and defensins (DA) in the nasopharyngeal milieu, during early phases of infection, and disease severity remains poorly understood. Therefore, we performed differential gene expression analysis of matrix metalloproteinases (MMPs) and defensins in the nasopharyngeal swab samples collected from mild and severe COVID-19 cases within three days of infection and examined the association between MMP and DA expression and disease severity.

**Material and Method:** A total of 118 previously collected nasopharyngeal samples from mild and severe COVID-19 patients (as per the WHO criteria) were used in this study. To determine the viral loads and assess the mRNA expression of matrix metalloproteinase (MMPs) and defensins, a real-time qPCR assay was used. To assess statistically significant differences in the mean expression of viral loads and the cytokines in between the severe and mild groups, an unpaired T-test was applied. The Pearson correlation test was used to assess the correlation between cytokine expressions. In addition, a multivariable logistic regression analysis was carried out with all the variables from the data set using ‘severity’ as the outcome variable.

**Results:** Our results showed that the expression of DA3 and MMP2 to be considerably lower in the severe group than in the mild group. Furthermore, there was a significant association between MMP1 and DA4 and DA6 (r=0.5, p=0.0001); as well as between MMP7 and DA1 and DA6 (r=0.5, p=0.00). Additionally, the regression analysis shows a significant correlation (p 0.05) between MMP2 and the severity of COVID-19 disease.

**Conclusion:** The early detection of changes in the expression of MMPs and defensins may act as a useful biomarker/predictor for possible severe COVID-19 disease, which may be useful in the clinical management of patients to reduce COVID-19-associated morbidity and mortality.

## Introduction

SARS-CoV-2, although significantly controlled by vaccine, remains a continuous threat to human health with substantial economic, social, and health implications [1-4]. Previous studies have shown that the SARS-CoV-2 virus has two possible entry routes, via endolysosomal cathepsins and the transmembrane serine protease 2 (TMPRSS2) [5].

The underlying pathophysiological mechanisms that are associated with the severe form of COVID-19 disease and death due to COVID-19-associated complications remain to be fully understood. The pre-existing comorbidities, e.g., diabetes, hypertension, compromised immune system, etc. have been associated with increased severity of COVID-19 [6, 7]. For example, the compromised immune system is associated with increased levels of metalloproteinases gelatinases, such as MMP-2 in plasma [8]. Therefore, it has been hypothesized that depending on the age and genetic polymorphisms, the pre-infection level of plasma matrix metalloproteinase (MMPs) or the potential of the host cells to secrete these proteases may be associated with the severe form of COVID-19 disease [9]. MMPs being proteolytic enzymes can damage various extracellular matrix components and are associated with the modulation of various cytokines and growth factors [10]. Recent data has also shown that increased plasma MMPs are associated with the severity of COVID-19 disease [11]. Furthermore, COVID-19-associated lung damage has also been shown to be associated with MMPs [12, 13], for instance, upregulation of gene expression of MMP-2 and MMP-9 in COVID-19 patients is associated with increased risk of respiratory failure [14, 15]. Additionally, is has been reported that MMPs can facilitate the viral entry and formation of syncytia in the context of dysregulated immune response and hyperinflammation in COVID-19 patients. Therefore, MMPs such as cathepsins and serine proteases can be potential therapeutic targets to treat severe COVID-19 patients.

Similar to MMPs, defensins (antimicrobial peptides), have been proposed as immunologic factors in the pathophysiology of COVID-19 disease and its severity [16, 17]. Defensins can expressed by mucosal epithelial cells as part of the innate immune response against the colonization of various pathogens [18]. Mild forms of COVID-19 disease may be associated with effective expression of defensins, as it has been shown that the activity of α- and β-defensins play a major role in controlling upper respiratory tract viral infections [19]. Recently, defensins have been explored as potential antiviral therapeutic agents against SARS-CoV-2, however, the exact relationship between various defensins and SARS-CoV-2 infection, especially during the early days of infection, has yet to be elucidated [20, 21].

Thus, a detailed analysis of the expression profile of human defensin genes may be useful to understanding the role of the viral infection patterns, innate immune response, and its subsequent association with the COVID-19 disease. Therefore, we performed differential gene expression analysis of matrix metalloproteinases (MMPs) and defensins in the nasopharyngeal swab samples collected from mild and severe COVID-19 cases within three days of infection and examined the association between MMP and DA expression and disease severity.

## Methodology

### Sample collection and characterization of the sample as mild and severe based on the patient’s symptoms

This was a retrospective, cross-sectional study, performed on a total of 118 SAR-CoV-2 PCR-positive nasopharyngeal swab samples. As described previously, using the WHO diagnostic criteria, these samples were characterized as severe and mild, based on symptoms observed in the patients [22-24]. These samples were obtained after written informed consent: the samples for the mild group were taken within three days of symptoms appealing, while samples for the severe group were taken within three days of admission to the hospital, and stored at -80°C until further use [24, 25]. The study was approved by the Ethics Review Committee, Aga Khan University Hospital (ERC#2021-5456-15382).

### RNA extraction, cDNA synthesis, estimation of viral loads, and gene expression analysis of MMP and defensin genes

Viral RNA was extracted from all the SARS-CoV-2 positive nasopharyngeal patient samples using a QIAamp viral RNA kit (Qiagen, Hilden, Germany). For reverse transcription, 2.5ug RNA/20ul of cDNA synthesis reaction was carried out using ONE SCRIPT PLUS cDNA Synthesis Kit (CAT # G236, ABM) as described previously [25]. The SARS-CoV-2 viral loads were accessed using COVID-19 genesis Real-Time PCR assay on CFX96 Touch Real-Time PCR System [25] using following thermocycling conditions: 55°C for 10 min, 95°C for 2 min followed by 45 cycles of 95°C for 10s, 60°C for the 60s. The Ct values of Internal control and Target (RdRp) genes were measured on Hex and FAM channels. The Ct values were used to assess viral load in each sample [25].

A qPCR assay was used to assess the gene expression profiles of MMPs and defensins in all samples employing gene-specific primers (Table 1), while beta-actin was used as a housekeeping gene and for normalization of the gene expression data. For qPCR reaction, 2ul of cDNA was mixed with 4ul of BlasTaq 2X qPCR Master mix (Cat # G891; ABM), and 0.5ul of each reverse and forward primers. The qPCR was run using the following thermal cycling conditions: 95°C for 3 minutes, 40 cycles of 95°C for 15 seconds, and 57.8°C to 64°C (depending on the primer) for 1 minute with a melt curve at 55-95_o_C. All reactions were run in duplicate. To compare/plot the expression of each cytokine in both groups, the delta Ct method was used, while to estimate the relative expression/fold change of each MMP and defensins in the severe vs mild group, 2_(-ΔΔCt)_ methods were used [26, 27].

**Table 1:**
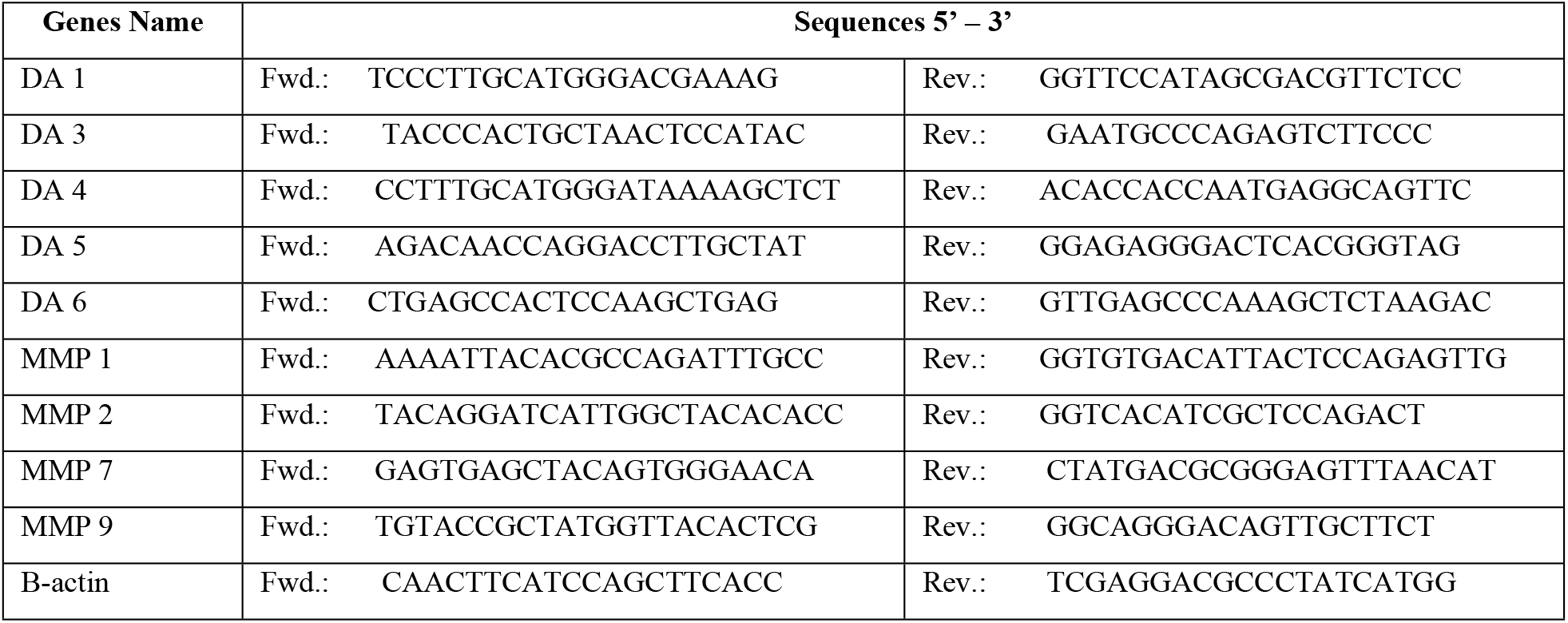
List of Primers used to measure the levels of matrix metalloproteinase and defensin and β-actin.

### Statistical Analyses

To assess statistically significant differences in the expression of viral loads and MMP and defensin genes, between the severe and mild groups, an unpaired T test was applied. Additionally, the Pearson correlation test was used to assess the correlation between cytokine expressions. In addition, a multivariable logistic regression analysis was carried out with all the variables from the data set using ‘severity’ as the outcome variable. For all statistical analyses, p*<*0.05 was considered significant. All statistical analyses were performed using the IBM SPSS software v20.

## Results

### Patient characteristics, viral load distribution in mild and severe groups, and correlation with disease severity

A total of 118 nasopharyngeal swab samples were used in this study; out of which 71 and 47 were from patients with mild and severe COVID-19 disease, as described previously [25]. Of these 47 patients with severe COVID-19 disease, 32 (68.0%) were male and 15 (31.25%) were female. The results of the descriptive statistics show that the mild group had significantly (p=0.002) lower age (mean = 44.1 years, SD = 18.03) than the severe group (mean = 54.61 years, SD = 17.26). The mean viral loads for the mild and severe groups were 27.07± 5.22 and 26.37±7.89, respectively; however, no association was found between viral load and disease severity (p>0.05).

### Analysis of differential expression of Defensins and MMPs in the mild and severe groups

The gene expression analysis showed the expression of only DA3 and MMP2 (severe: 26.53±3.358 STDEV, mild: 23.56±2.740 STDEV; p<0.0001) to be significantly different in mild versus severe groups (Figure 1A). The fold change analysis showed the expression of DA1, DA3, DA4, DA5, and DA6 to be, respectively, 1.82-fold lower, 3.90-fold lower, 6.39-fold higher, 1.33-fold lower, 3.02-fold lower expression in the severe group as compared to the mild group. Similarly, the fold change analysis of MMPs showed the expression of MMP1, MMP2, MMP7, and MMP9 to be, respectively, 2.48-fold higher, 7.71-fold lower, 1.04-fold higher, 1.64-fold lower, in the severe group as compared to the mild group (Figure 1B).

**Figure 1.**
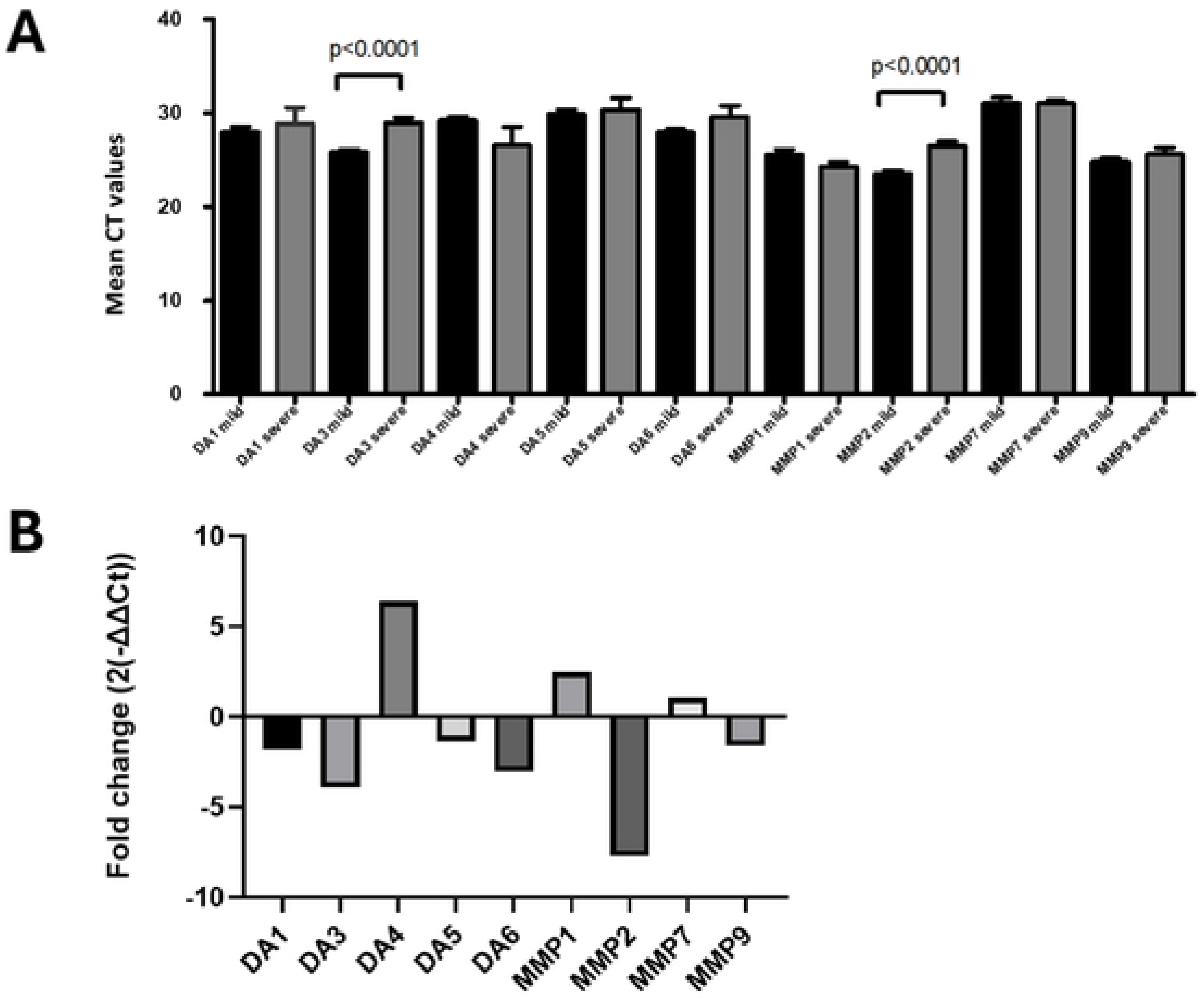
Relative expression of defensins (DAs) and matrix metalloproteinases (MMPs) in mild vs severe group. The figure shows the **A**) mean ΔCt values and **B**) fold change in the mild group as compared to the severe group. **A)** The line above bars indicate a significant difference (p*<*0.05) in the expression of the tested genes between the mild and severe groups.

### Correlation between Defensins and Matrix Metalloproteinases expression in mild and severe groups

In the next step, the Pearson correlation test was applied to investigate the relationship between the expression of defensins and matrix metalloproteinase in mild and severe groups (Table 2).

**Table 2:** Correlation among Defensins and Matrix Metalloproteinase expressions in the mild group. Statistically significant values are highlighted in bold.

| Mild group |  |  |  |  |  |
| --- | --- | --- | --- | --- | --- |
| Gene Expression | DA1 | DA3 | DA4 | DA5 | DA6 |
| DA1 | - | **r= 0.5** | **r= 0.5** | **r=0.5** | **r=0.60** |
| DA3 | r= 0.5 | - | **r =0.6** | **r=0.5** | **r=0.7** |
| DA4 | r= 0.5 | r =0.6 | - | **r=0.5** | **r =0.6** |
| DA5 | r= 0.5 | r= 0.5 | r= 0.5 | - | r= 0.8 |
| DA6 | r =0.6 | r=0.7 | r =0.6 | **r= 0.8** | - |
| MMP1 | r= 0.2 | **r= 0.3** | **r= 0.4** | r= 0.00 | r= 0.1 |
| MMP2 | r= 0.2 | r= 0.2 | **r= 0.3** | r= 0.2 | r= 0.00 |
| MMP7 | **r= 0.3** | r= 0.00 | **r= 0.3** | **r= 0.3** | r= 0.2 |
| MMP9 | r= 0.1 | r= 0.2 | r= 0.2 | r= 0.2 | **r= 0.4** |
|  | **MMP1** | **MMP2** | **MMP7** | **MMP9** |  |
| MMP1 | - | r=0.2 | r=0.1 | r=0.0 |  |
| MMP2 | r=0.2 | - | r=0.0 | **r=0.4** |  |
| MMP7 | r=0.1 | r=0.0 | - | r=0.0 |  |
| MMP9 | r=0.0 | r=0.4 | r=0.0 | - |  |
| Severe group |  |  |  |  |  |

| Gene Expression | DA1 | DA3 | DA4 | DA5 | DA6 |
| --- | --- | --- | --- | --- | --- |
| DA1 | - | **r= 0.9** | r= 0.1 | r=0.2 | **r=0.7** |
| DA3 | r= 0.9 | - | r=0.1 | r=0.1 | **r=0.5** |
| DA4 | r= 0.1 | r= 0.1 | - | **r= 0.4** | r= 0.0 |
| DA5 | r= 0.2 | r= 0.1 | r= 0.4 | - | r= 0.1 |
| DA6 | r= 0.7 | r= 0.5 | r= 0.0 | r= 0.1 | - |
| MMP1 | **r= 0.4** | r= 0.2 | **r= 0.4** | r= 0.5 | **r= 0.5** |
| MMP2 | r= 0.3 | r= 0.00 | r= 0.3 | **r= 0.4** | **r= 0.4** |
| MMP7 | **r= 0.5** | r= 0.2 | **r= 0.4** | **r= 0.4** | **r= 0.5** |
| MMP9 | r= 0.2 | r= 0.1 | r= 0.2 | r= 0.2 | r= 0.1 |
|  | **MMP1** | **MMP2** | **MMP7** | **MMP9** |  |
| MMP1 | - | **r= 0.6** | **r= 0.9** | **r= 0.3** |  |
| MMP2 | r= 0.6 | - | **r= 0.6** | r= 0.0 |  |
| MMP7 | r= 0.9 | **r= 0.6** | - | r= 0.4 |  |
| MMP9 | r= 0.3 | r= 0.0 | **r= 0.4** | - |  |

In the mild group, a statistically significant, but moderate positive correlation was found between DA3 and MMP1 (r= 0.03, p=0.02), DA1 and MMP7 (r=0.3, p=0.01), and DA4 with the matrix metalloproteinase MMP1 (r=0.3, p=0.001), MMP2 (r=0.03, p=0.001), MMP7 (r=0.3, p =0.001), DA5 and MMP7 (r=0.3, p =0.001), DA6 and MMP9 (r=0.4, p =0.001) (Table 2). While amongst defensins, a statistically significant strong correlation was observed amongst the defensins DA1 and DA3 (r =0.5, p=0.001), DA1 and DA4 (r=0.5, p=0.001), DA1 and DA5 (r=0.5, p=0.001), DA1 and DA6 (r=0.6, p=0.001), DA3 and DA4 (r= 0.6, p=0.001), DA3 and DA5 (r=0.5, p=0.01), DA3 and DA6 (r=0.7, p=0.001), DA4 and DA5 (r=0.5, p=0.001), DA4 and DA6 (r=0.6, p=0.001), DA5 and DA6 (r=0.8, p=0.001) (Table 4b). Moreover, a statistically significant strong positive correlation was only found between MMP2 and MMP9 (r=0.4, p=0.001).

In the severe group, there was a statistically significant and strong positive correlation between MMP1 and the defensins DA6 (r= 0.5, p=0.001) and DA4 (r= 0.5, p= 0.001), and MMP7 with the defensins DA1 (r= 0.5, p= 0.001) and DA6 (r= 0.5, p= 0.001). Further, a statistically strong but moderate correlation was observed between MMP1 with DA1 (r= 0.4, p= 0.001), MMP7 with DA1 (r= 0.4, p= 0.001), DA5 (r= 0.4, p= 0.001) and DA4 (r= 0.4, p= 0.001), and MMP2 with DA5 and DA6 (r= 0.4, p= 0.001). Amongst defensins, a statistically significant and strong positive correlation between DA1 and DA3 (r =0.9, p=0.001), DA1 and DA6 (r=0.7, p=0.001), DA3 and DA6 (r=0.5, p=0.001), DA4 and DA5 (r=0.4, p=0.001) was also observed (Table 2). Moreover, among MMP genes statistically strong positive correlation was found only between MMP1 and MMP2 (r=0.6, p=0.001), MMP1 and MMP7 (r=0.9, p=0.001), MMP1 and MMP9 (r=0.3, p=0.001), MMP2 and MMP7 (r=0.6, p=0.001), MMP7 and MMP9 (r=0.4, p=0.001).

### Multivariate regression analysis

Regression analysis was performed to examine the influence of all study variables with outcome ‘severity’. The results showed that the model as a whole was significant (Chi_2_(12) = 53.04, p <0.001, n = 117). Regression analysis also showed only age (p=0.003), and expression of MMP1, and MMP2 (p<0.001) to be significantly associated with severity (Table 3). The odds ratio of 1.03 for the variable age suggests that a one-unit increase in the variable age will increase the odds of getting the severe disease by 1.03 times. Similarly, the odds ratio of 1.68 for MMP2 suggests that a one-unit increase in MMP2 expression will increase the odds of severe disease by 1.68 times (Table 4). The odds ratio of 0.84 for MMP1 suggests that a one-unit increase in MMP1 expression will decrease the odds of severe disease by 0.84 times (Table 3).

**Table 3.** Multivariate logistic regression model in different variables.

|  | Coefficient<br>B | Standard<br>error | z | p | Odds<br>Ratio | 95% conf.<br>interval |
| --- | --- | --- | --- | --- | --- | --- |
| Age | **0.03** | **0.02** | **1.67** | **0.003** | **1.03** | **1 - 1.06** |
| Gender | -0.48 | 0.57 | 0.86 | 0.393 | 0.62 | 0.2 - 1.87 |
| Viral Ct<br>value | 0.05 | 0.05 | 1.07 | 0.283 | 1.05 | 0.96 - 1.15 |
| DA1 | 0.09 | 0.09 | 0.98 | 0.33 | 1.09 | 0.91 - 1.31 |
| DA3 | 0.15 | 0.08 | 1.84 | 0.066 | 1.16 | 0.99 - 1.37 |
| DA4 | -0.08 | 0.04 | 1.88 | 0.06 | 0.92 | 0.85 - 1 |
| DA5 | 0.06 | 0.05 | 1.09 | 0.275 | 1.06 | 0.95 - 1.18 |
| DA6 | 0.08 | 0.06 | 1.19 | 0.235 | 1.08 | 0.95 - 1.23 |
| **MMP1** | **-0.17** | **0.08** | **2.21** | **0.027** | **0.84** | **0.73 - 0.98** |
| **MMP2** | **0.52** | **0.14** | **3.65** | **<.001** | **1.68** | **1.27 - 2.22** |
| MMP7 | 0.03 | 0.06 | 0.44 | 0.659 | 1.03 | 0.91 - 1.16 |
| MMP9 | 0.03 | 0.04 | 0.71 | 0.479 | 1.03 | 0.95 - 1.11 |

## DISCUSSION

Studies have shown that the COVID-19 disease course may vary from mild respiratory disease to severe disease with associated complications and high mortality [28]. The severity of the disease may be affected by several factors such as the viral load, age, gender, and dysregulated expression of antiviral/proinflammatory cytokines [25, 29].

In our study, the expression of all tested defensins, except defensin 3, defensin 4, and defensin 5 was found to be comparable between the two groups. Only defensin 3 and defensin 5 showed, respectively, 3.9 and 3-fold lesser expression, while defensin 4 showed 6-fold higher expression in the severe group as compared to the mild group. Overall, defensins exhibit some degree of antiviral properties, preventing the virus from entering the cell, thereby, inhibiting infection of the virus [30]. However, it is still unclear how each specific defensins affect the COVID-19 severity. Therefore, human defensins and their antiviral role are one of the major active areas of investigation [31], with studies showing defensins 4/2 to be dysregulated in COVID-19 patients [17]. Defensin 5 has also been shown to impede the entry of SARS-CoV-2 into human renal proximal tubular epithelial cells, potentially mitigating the severity of COVID-19 [32]. Studies have reported the possible inhibition of SARS-CoV-2 spike protein-mediated fusion by the defensins, for instance, HNP1 was reported to have weak inhibition of the virus attachment [33]. Furthermore, virus-mediated decreased expression of various defensin genes in COVID-19 patients may result in enhanced added bacterial infections in the upper respiratory tract; therefore, agents that can enhance the expression of human defensin genes (HBD-1-3) may have a potential therapeutic effect in these patients [34, 35].

In our study, the expression of all tested MMPs, except MMP1 and MMP2, was found to be comparable between the two groups. MMP1 showed 2.48-fold higher, while MMP2 showed 7.7-fold lower expression in the severe group compared to the mild group, further confirmed by logistic regression analysis. The role of the plasma levels of MMP2 has been reported in hypertensive COVID-19 patients [36]. There are debates about which MMPs cause the more severe COVID-19 disease. Some studies state that MMP1 impacts the disease severity by damaging extracellular matrix (ECM) components that lead to tissue damage and inflammation [37, 38]. On the other hand, other studies found that MMP2 is associated with severe COVID-19 due to hyperinflammation and lung tissue damage caused by the decrease in collagen levels [39]. Higher expression of MMP2 has been found in the tracheal-aspirate fluid samples of patients with severe COVID-19 patients [40], however, we found decreased expression of MMP2 in the nasopharyngeal milieu in patients with severe COVID-19, which may suggest differences in the expression of MMPs in different anatomical location and/or cells [41, 42]. Studies have shown that SARS-CoV-2 and SARS-CoV-1 use ACE2-dependent pathways involving an endosomal cathepsin protease pathway and a surface serine protease pathway [43, 44]. In virus-producing cells, the proteolytic processing at the S1/S2 boundary required a higher expression of MMP9 and MMP2 proteases, which is associated with hyperinflammation and lung tissue damage in patients with COVID-19 patients [45].

A significant correlation between the age of patients and SARS-CoV-2 infection was observed, which aligns with previous research [46]. The severity of COVID-19 is influenced by age: younger infected patients had milder disease due to protective mechanisms. In contrast, older patients rely more on memory T cells for immune response, potentially leading to overreaction and tissue damage, which makes the disease more severe [29].

We acknowledge certain limitations of our study. Firstly, the sample size was small to establish predictors of severity, however, we believe it was sufficient to show the direct correlation between the disease severity and defensins and MMP-2 expression, which also agrees with the results of previous studies. Secondly, the other possible reasons for COVID-19 severity could not be established due to the non-availability of clinical information about other co-existing medical conditions that may be present in the study participants. Finally, the expression of MMPs was analyzed only in the nasopharyngeal swab samples and not in the serums, which may result in different expression profiles.

In conclusion, we found altered nasopharyngeal expression of certain defensins and MMPs (MMP-2) in the severe group. These findings demonstrate that detection of dysregulated expression of defensins and MMPs in the nasopharyngeal milieu, which can be observed as early as three days (time of our sample collection) might be correlated with the severe form of the disease. Therefore, the early estimation of the expression of these genes may act as a useful biomarker/predictor for possible severe COVID-19 disease, which may be useful in the clinical management of patients to reduce COVID-19-associated morbidity and mortality.

## Acknowledgment

We are grateful to Dr Abid Jamal, Cancer Foundation Hospital, Karachi, Pakistan, for providing access to the samples.

## Author Contributions

Conceptualization: SHA; Methodology and formal analysis: KI, NF, KA, MFA, AH, ASS, KG; Writing – original draft: KI, NF, KA, AZ; Review and Final draft: SHA. Supervision: SHA. Funding acquisition: SHA.

## References

1. Hardy E, Fernandez-Patron C. Targeting MMP-Regulation of Inflammation to Increase Metabolic Tolerance to COVID-19 Pathologies: A Hypothesis. Biomolecules. 2021;11(3). Epub 2021/04/04. doi: 10.3390/biom11030390. PubMed PMID: 33800947; PubMed Central PMCID: PMCPMC7998259.

2. Ayres JS. Surviving COVID-19: A disease tolerance perspective. Sci Adv. 2020;6(18):eabc1518. Epub 2020/06/05. doi: 10.1126/sciadv.abc1518. PubMed PMID: 32494691; PubMed Central PMCID: PMCPMC7190329.

3. Ayres JS. A metabolic handbook for the COVID-19 pandemic. Nat Metab. 2020;2(7):572–85. Epub 2020/07/23. doi: 10.1038/s42255-020-0237-2. PubMed PMID: 32694793; PubMed Central PMCID: PMCPMC7325641.

4. Schneider DS, Ayres JS. Two ways to survive infection: what resistance and tolerance can teach us about treating infectious diseases. Nat Rev Immunol. 2008;8(11):889–95. Epub 2008/10/18. doi: 10.1038/nri2432. PubMed PMID: 18927577; PubMed Central PMCID: PMCPMC4368196.

5. Murgolo N, Therien AG, Howell B, Klein D, Koeplinger K, Lieberman LA, et al. SARS-CoV-2 tropism, entry, replication, and propagation: Considerations for drug discovery and development. PLoS Pathog. 2021;17(2):e1009225. Epub 2021/02/18. doi: 10.1371/journal.ppat.1009225. PubMed PMID: 33596266; PubMed Central PMCID: PMCPMC7888651 Co., Inc., Kenilworth, NJ, USA and may own stock or hold stock options in Merck & Co., Inc., Kenilworth, NJ, USA.

6. Guan WJ, Ni ZY, Hu Y, Liang WH, Ou CQ, He JX, et al. Clinical Characteristics of Coronavirus Disease 2019 in China. N Engl J Med. 2020;382(18):1708–20. Epub 2020/02/29. doi: 10.1056/NEJMoa2002032. PubMed PMID: 32109013; PubMed Central PMCID: PMCPMC7092819.

7. Xie J, Wu W, Li S, Hu Y, Hu M, Li J, et al. Clinical characteristics and outcomes of critically ill patients with novel coronavirus infectious disease (COVID-19) in China: a retrospective multicenter study. Intensive Care Med. 2020;46(10):1863–72. Epub 2020/08/21. doi: 10.1007/s00134-020-06211-2. PubMed PMID: 32816098; PubMed Central PMCID: PMCPMC7439240.

8. Malemud CJ. Matrix metalloproteinases (MMPs) in health and disease: an overview. Front Biosci. 2006;11:1696–701. Epub 2005/12/22. doi: 10.2741/1915. PubMed PMID: 16368548.

9. Salomão R, Assis V, de Sousa Neto IV, Petriz B, Babault N, Durigan JLQ, et al. Involvement of matrix metalloproteinases in COVID-19: molecular targets, mechanisms, and insights for therapeutic interventions. Biology. 2023;12(6):843.

10. Fernandez-Patron C, Kassiri Z, Leung D. Modulation of Systemic Metabolism by MMP-2: From MMP-2 Deficiency in Mice to MMP-2 Deficiency in Patients. Compr Physiol. 2016;6(4):1935–49. Epub 2016/10/27. doi: 10.1002/cphy.c160010. PubMed PMID: 27783864.

11. C Da-M, Couto AES, Campos LCB, Vasconcelos TF, Michelon-Barbosa J, Corsi CAC, et al. MMP-2 and MMP-9 levels in plasma are altered and associated with mortality in COVID-19 patients. Biomed Pharmacother. 2021;142:112067. Epub 2021/08/28. doi: 10.1016/j.biopha.2021.112067. PubMed PMID: 34449310; PubMed Central PMCID: PMCPMC8376652.

12. Davey A, McAuley DF, O’Kane CM. Matrix metalloproteinases in acute lung injury: mediators of injury and drivers of repair. Eur Respir J. 2011;38(4):959–70. Epub 2011/05/14. doi: 10.1183/09031936.00032111. PubMed PMID: 21565917.

13. Fligiel SE, Standiford T, Fligiel HM, Tashkin D, Strieter RM, Warner RL, et al. Matrix metalloproteinases and matrix metalloproteinase inhibitors in acute lung injury. Hum Pathol. 2006;37(4):422–30. Epub 2006/03/28. doi: 10.1016/j.humpath.2005.11.023. PubMed PMID: 16564916.

14. Hazra S, Chaudhuri AG, Tiwary BK, Chakrabarti N. Matrix metallopeptidase 9 as a host protein target of chloroquine and melatonin for immunoregulation in COVID-19: A network-based meta-analysis. Life Sci. 2020;257:118096. Epub 2020/07/18. doi: 10.1016/j.lfs.2020.118096. PubMed PMID: 32679150; PubMed Central PMCID: PMCPMC7361122.

15. Ueland T, Holter JC, Holten AR, Muller KE, Lind A, Bekken GK, et al. Distinct and early increase in circulating MMP-9 in COVID-19 patients with respiratory failure. J Infect. 2020;81(3):e41–e3. Epub 2020/07/01. doi: 10.1016/j.jinf.2020.06.061. PubMed PMID: 32603675; PubMed Central PMCID: PMCPMC7320854.

16. Cao X. COVID-19: immunopathology and its implications for therapy. Nat Rev Immunol. 2020;20(5):269–70. Epub 2020/04/11. doi: 10.1038/s41577-020-0308-3. PubMed PMID: 32273594; PubMed Central PMCID: PMCPMC7143200.

17. Al-Bayatee NT, Ad’hiah AH. Human beta-defensins 2 and 4 are dysregulated in patients with coronavirus disease 19. Microbial Pathogenesis. 2021;160:105205.

18. Laneri S, Brancaccio M, Mennitti C, De Biasi MG, Pero ME, Pisanelli G, et al. Antimicrobial peptides and physical activity: a great hope against COVID 19. Microorganisms. 2021;9(7):1415.

19. Banu S, Nagaraj R, Idris MM. Defensins: Therapeutic molecules with potential to treat SARS-CoV-2 infection. Indian Journal of Medical Research. 2022;155(1):83–5.

20. Mabrouk DM. Antimicrobial peptides: features, applications and the potential use against covid-19. Molecular Biology Reports. 2022;49(10):10039–50.

21. Solanki SS, Singh P, Kashyap P, Sansi MS, Ali SA. Promising role of defensins peptides as therapeutics to combat against viral infection. Microbial pathogenesis. 2021;155:104930.

22. Son KB, Lee TJ, Hwang SS. Disease severity classification and COVID-19 outcomes, Republic of Korea. Bull World Health Organ. 2021;99(1):62–6. Epub 2021/03/05. doi: 10.2471/BLT.20.257758. PubMed PMID: 33658735; PubMed Central PMCID: PMCPMC7924894.

23. Ghulam U, Nazim F, Farooqui N, Rizwan-ul-Hasan S, Anwar MF, Ahmed K, et al. Analysis of differential gene expression of pro-inflammatory cytokines in the nasopharyngeal milieu of mild & severe COVID-19 cases. Plos one. 2022;17(12):e0279270.

24. Zahid W, Farooqui N, Zahid N, Ahmed K, Anwar MF, Rizwan-ul-Hasan S, et al. Association of Interferon Lambda 3 and 4 Gene SNPs and Their Expression with COVID-19 Disease Severity: A Cross-Sectional Study. Infection and Drug Resistance. 2023:6619–28.

25. Ghulam U, Nazim F, Farooqui N, Rizwan-Ul-Hasan S, Anwar MF, Ahmed K, et al. Analysis of differential gene expression of pro-inflammatory cytokines in the nasopharyngeal milieu of mild & severe COVID-19 cases. PLoS One. 2022;17(12):e0279270. Epub 2022/12/31. doi: 10.1371/journal.pone.0279270. PubMed PMID: 36584119; PubMed Central PMCID: PMCPMC9803207.

26. Schmittgen TD, Livak KJ. Analyzing real-time PCR data by the comparative C(T) method. Nat Protoc. 2008;3(6):1101–8. Epub 2008/06/13. doi: 10.1038/nprot.2008.73. PubMed PMID: 18546601.

27. Livak KJ, Schmittgen TD. Analysis of relative gene expression data using real-time quantitative PCR and the 2(-Delta Delta C(T)) Method. Methods. 2001;25(4):402–8. Epub 2002/02/16. doi: 10.1006/meth.2001.1262. PubMed PMID: 11846609.

28. Gisondi P, S Pi, Bordin C, Alaibac M, Girolomoni G, Naldi L. Cutaneous manifestations of SARS-CoV-2 infection: a clinical update. J Eur Acad Dermatol Venereol. 2020;34(11):2499–504. Epub 2020/06/26. doi: 10.1111/jdv.16774. PubMed PMID: 32585074; PubMed Central PMCID: PMCPMC7362144.

29. Brodin P. Immune determinants of COVID-19 disease presentation and severity. Nat Med. 2021;27(1):28–33. Epub 2021/01/15. doi: 10.1038/s41591-020-01202-8. PubMed PMID: 33442016.

30. Xu C, Wang A, Marin M, Honnen W, Ramasamy S, Porter E, et al. Human defensins inhibit SARS-CoV-2 infection by blocking viral entry. Viruses. 2021;13(7):1246.

31. Ahmed A, Siman-Tov G, Hall G, Bhalla N, Narayanan A. Human Antimicrobial Peptides as Therapeutics for Viral Infections. Viruses. 2019;11(8). Epub 2019/08/04. doi: 10.3390/v11080704. PubMed PMID: 31374901; PubMed Central PMCID: PMCPMC6722670.

32. Wang C, Wang, S., Li, D., Wei, D. Q., Zhao, J., & Wang, J. Human Intestinal Defensin 5 Inhibits SARS-CoV-2 Invasion by Cloaking ACE2. Gastroenterology. 2020;159(3):1145–7. doi: 10.1053/j.gastro.2020.05.015.

33. Xu D, Lu W. Defensins: A Double-Edged Sword in Host Immunity. Front Immunol. 2020;11:764. Epub 2020/05/28. doi: 10.3389/fimmu.2020.00764. PubMed PMID: 32457744; PubMed Central PMCID: PMCPMC7224315.

34. Krishnakumari V, Guru A, Adicherla H, Nagaraj R. Effects of increasing hydrophobicity by N-terminal myristoylation on the antibacterial and hemolytic activities of the C-terminal cationic segments of human-beta-defensins 1-3. Chem Biol Drug Des. 2018;92(2):1504–13. Epub 2018/04/24. doi: 10.1111/cbdd.13317. PubMed PMID: 29682907.

35. Hoover DM, Wu Z, Tucker K, Lu W, Lubkowski J. Antimicrobial characterization of human beta-defensin 3 derivatives. Antimicrob Agents Chemother. 2003;47(9):2804–9. Epub 2003/08/26. doi: 10.1128/AAC.47.9.2804-2809.2003. PubMed PMID: 12936977; PubMed Central PMCID: PMCPMC182640.

36. Carolina D, Couto AE, Campos LC, Vasconcelos TF, Michelon-Barbosa J, Corsi CA, et al. MMP-2 and MMP-9 levels in plasma are altered and associated with mortality in COVID-19 patients. Biomedicine & Pharmacotherapy. 2021;142:112067.

37. Herrera I CJ, Maldonado M, Ramírez R, Ortiz-Quintero B, Anso E, Chandel NS, Selman M, Pardo Matrix metalloproteinase (MMP)-1 induces lung alveolar epithelial cell migration and proliferation, protects from apoptosis, and represses mitochondrial oxygen consumption. J Biol Chem. 2013;288(36):25964–75. doi: 10.1074/jbc.M113.459784.

38. Choreño-Parra JA J-ÁL, Cruz-Lagunas A, Rodríguez-Reyna TS, Ramírez-Martínez G, Sandoval-Vega M, Hernández-García DL, Choreño-Parra EM, Balderas-Martínez YI, Martinez-Sánchez ME, Márquez-García E, Sciutto E, Moreno-Rodríguez J, Barreto-Rodríguez JO, Vázquez-Rojas H, Centeno-Sáenz GI, Alvarado-Peña N, Salinas-Lara C, Sánchez-Garibay C, Galeana-Cadena D, Hernández G, Mendoza-Milla C, Domínguez A, Granados J, Mena-Hernández L, Pérez-Buenfil L, Domínguez-Cheritt G, Cabello-Gutiérrez C, Luna-Rivero C, Salas-Hernández J, Santillán-Doherty P, Regalado J, Hernández-Martínez A, Orozco L, Ávila-Moreno F, García-Latorre EA, Hernández-Cárdenas CM, Khader SA, Zlotnik A, Zúñiga J. Clinical and immunological factors that distinguish COVID-19 from pandemic influenza A (H1N1). Front Immunol. 2021;12:593595. doi: 10.3389/fimmu.2021.593595.

39. Gustavo R.M. LAJ, Alfredo C.L., Sergio I.C., Itzel A.G., Tatiana S.R., Jose A.C., & Joaquí Z. Possible Role of Matrix Metalloproteinases and TGF-β in COVID-19 Severity and Sequelae. Journal of Interferon & Cytokine Research. 2022;42(8):352–68. doi: 10.1089/jir.2021.0222. PubMed PMID: 35647937.

40. da Silva-Neto PV, do Valle VB, Fuzo CA, Fernandes TM, Toro DM, Fraga-Silva TF, et al. Matrix metalloproteinases on severe COVID-19 lung disease pathogenesis: cooperative actions of MMP-8/MMP-2 axis on immune response through HLA-G shedding and oxidative stress. Biomolecules. 2022;12(5):604.

41. Miller TL, Touch SM, Shaffer TH. Matrix metalloproteinase and tissue inhibitor of matrix metalloproteinase expression profiles in tracheal aspirates do not adequately reflect tracheal or lung tissue profiles in neonatal respiratory distress: observations from an animal model. Pediatr Crit Care Med. 2006;7(1):63–9. doi: 10.1097/01.pcc.0000192320.87416.1a. PubMed PMID: 16395077.

42. Brand KH, Ahout IM, de Groot R, Warris A, Ferwerda G, Hermans PW. Use of MMP-8 and MMP-9 to assess disease severity in children with viral lower respiratory tract infections. J Med Virol. 2012;84(9):1471–80. doi: 10.1002/jmv.23301. PubMed PMID: 22825827; PubMed Central PMCID: PMCPMC7167016.

43. Shang J, Wan Y, Luo C, Ye G, Geng Q, Auerbach A, et al. Cell entry mechanisms of SARS-CoV-2. Proc Natl Acad Sci U S A. 2020;117(21):11727–34. Epub 2020/05/08. doi: 10.1073/pnas.2003138117. PubMed PMID: 32376634; PubMed Central PMCID: PMCPMC7260975.

44. Hoffmann M, Hofmann-Winkler H, Smith JC, Kruger N, Arora P, Sorensen LK, et al. Camostat mesylate inhibits SARS-CoV-2 activation by TMPRSS2-related proteases and its metabolite GBPA exerts antiviral activity. EBioMedicine. 2021;65:103255. Epub 2021/03/08. doi: 10.1016/j.ebiom.2021.103255. PubMed PMID: 33676899; PubMed Central PMCID: PMCPMC7930809.

45. Benlarbi M, Laroche G, Fink C, Fu K, Mulloy RP, Phan A, et al. Identification of a SARS-CoV-2 host metalloproteinase-dependent entry pathway differentially used by SARS-CoV-2 and variants of concern Alpha, Delta, and Omicron. bioRxiv. 2022:2022.02.19.481107. doi: 10.1101/2022.02.19.481107.

46. Kim M-H SS, Wang JY, et al. Type I, II, and III interferon signatures correspond to coronavirus disease 2019 severity. J Infect Dis. 2021;224(5):777–82. doi: 10.1093/infdis/jiab288.

